# Thermodynamic stability of G-quadruplex and overlapping m6A: Position-dependent collaborators in allele frequency and fitness?

**DOI:** 10.1101/2021.04.28.441767

**Authors:** Cagri Gulec

## Abstract

**Background:** Post-transcriptional modifications like m6A, and secondary structures like G-quadruplex (G4), play an important role in RNA processing. Despite an emerging number of studies focusing on m6A and G4 separately, there are less studies about their synergy.

**Aim:** Since m6A is known to be enzymatically created in DRACH-motif, and genetic variants may create a novel DRACH-motif or abolish a pre-existing DRACH-motif, we can suppose that the variants may affect gene product level through modulating m6A-G4 colocalization, which consequently may affect fitness and change allele frequency. To test this hypothesis, rare and common variants in selected human genes were investigated in terms of their effect on m6A-G4 colocalization.

**Methods:** Genomic sequences and variant features were fetched from GRCh37/hg19 and Biomart-Ensembl databases, respectively. Counting the number of putative m6A- and G4-motifs in sequences and statistical analysis were performed with appropriate libraries of Python3.7.

**Results:** Common variants creating novel m6A-motif were found more frequently inside than outside G4, and displayed unequal distribution throughout pre-mRNA. Unequal distribution of m6A-creating variants seemed to be related to their effect on thermodynamic stability of the overlapping-G4.

**Discussion:** Selective m6A-G4 colocalization suggests that m6A-motif is favorable when overlapping with G4. Besides, thermodynamic stability may lead to unequal distribution of m6A-G4 colocalization, because m6A-creating alleles seem to have lower frequency if stabilizes overlapping-G4 in 3-prime-side, but not in 5-prime-side. We can conclude that the fitness, and consequently frequency of an m6A-creating variant is prone to become higher or lower depending on its position and effect on the overlapping-G4 stability.

## Intruduction

Genetic variants in protein-coding genes display their impact on the phenotype mostly through quantitative and qualitative fluctuation in protein products. In this respect, post-transcriptional RNA modifications play an important role especially in quantitative manner. To date, more than 150 chemical modifications have been identified in RNAs [1, 2]. Though most of these modifications were found in non-coding RNAs [rRNA, tRNA, snRNA and long-non-coding RNs), it has been shown that coding RNAs are also subject to such modifications [3, 4]. N6-methyladenosine (m6A) is one of the most abundant chemical modifications found in mRNAs [5, 6]. The function of m6A began to be elucidated long after its discovery [7, 8]. We now know that m6A plays an important role in processing, nuclear export, translational regulation and decay of the mRNAs [9, 10]. In line with these functions, m6A modification was found to display an unequal distribution throughout mRNA [5, 6, 11, 12].

In addition to post-transcriptional modifications, secondary structures have also important role in translational efficiency of the mRNAs. One of these secondary structures is the G-quadruplex (G4) that is spontaneously formed in guanine-rich (G-rich) sequences in DNA or RNA. This structure is known to be formed via Hoogsten base pairing of adjacent guanines in G-rich sequences [13]. Although Hoogsten hydrogen binds between G bases is essential for G4 structure, there are many additional factors that may affect G4 folding in a G-rich sequence, like the length of the sequence, the presence of an alternative Watson-Crick pair-based stable structure and free metal cations [14–17]. Both bioinformatic studies based on the canonical consensus sequence, and experimental genome-wide studies using G4 specific probes have revealed a non-random distribution of G4 structure throughout the genome and transcriptome [18]. These studies demonstrated that the G4 sequences are enriched at telomeres, promoter regions and replication origins in genomic DNA, and UTRs in mRNA. Though G4 structures are formed both in DNA and RNA, it has been shown that RNA G4 structures are more stable and have less topological diversity than G4 structures in DNA [19]. Searching the canonical consensus sequence (5⍰-G_3_N_1–7_G_3_N_1–7_G_3_N_1–7_G_3_-3⍰ where N is A, C, G, T, or U) from which a typical G4 structure is known to form [17, 20, 21], putative G4 structures were found to be present in 5′UTR of more than 9.000, and in 3′UTR of more than 8.000 human genes [22–24]. Moreover, it was shown that more than 1600 human genes have G4 structures in their ORF [25]. G4 structures in RNAs are supposed to play role in stability, splicing and translation of RNAs through binding specific proteins like eIF4G, LARK, SLIRP, AFF3, AFF4, eIF4A and hnRNP A2 [26–32].

Despite the great difference in their formation and structure, m6A modification and G4 structure seem to share common functional consequences in RNA processing. Both were separately shown to modulate splicing, nuclear export, translation and decay of the RNAs. Although, mutual or synergic activities of m6A and G4 were not well understood, recent studies have revealed that m6A modification can modulate G4 structure formation and vice versa. The m6A modification was found to modulate G4 structure through affecting its stability as shown in R-loops [33], while the G4 structure was demonstrated to modulate m6A modification through facilitating the adenine N6-methylation in target motif as shown in viral RNAs [34]. Their synergic activity was supported by overlapping m6A-G4 in eukaryotic mRNAs, as well [35, 36].

Taking into account the previous studies, this study aimed to investigate whether the functional consequences of overlapping m6A-G4 may have selective pressure on the variants that lead colocalization of m6A and G4. To address this question, defined single nucleotide variants in selected disease-associated human genes were investigated in terms of their effect on the colocalization of m6A and G4.

## Methods

### Reference sequence data

Human genome sequence (GRCh37/hg19) was downloaded from NCBI (https://www.ncbi.nlm.nih.gov/assembly/GCF_000001405.25) and recorded as a sql database using Sqlite3 library of Python3.7. Name list, exonic coordinates and variant coordinates of disease associated human genes (28,250 transcripts from 3,306 MIM genes) were fetched from Ensembl (Biomart; http://grch37.ensembl.org/biomart/martview/c4f6ef5ffc88cf1d9a00b63173228cda) and recorded as text files.

### Artificial variant sequences

Considering their genomic position, strand, alleles and allele frequency, each single nucleotide variant and their flanking reference sequence was fetched from hg19-sql database, and separately stored as ‘wild-type sequence pieces’. By replacing reference base with variant base, a ‘variant sequence piece’ was produced for each corresponding ‘wild-type sequence piece’. Each ‘variant sequence piece was tagged as a ‘common’ or ‘rare’ regarding the minor allele frequency, MAF (‘common’ for alleles with MAF >=0.01, and ‘rare’ for alleles with MAF <0.01) of the corresponding variant. Variants with no recorded MAF value in database were evaluated as rare variant.

### Searching and counting the putative G4 structures and m6A motifs

Both wild-type and matched variant sequence pieces were evaluated with ‘re’ module and ‘Pandas’ library of Python3.7, in terms of the presence and the count of putative G4-forming sequence (G_1-3_[N_1-7_G_1-3_]_3_) for putative G4 structures and DRACH motif ([A/G/U][A/G]AC[A/C/U]) for putative m6A modification sites. The effect of the variant on the m6A motif number was evaluated according to m6A motif Number (n) in the presence of reference allele. Resulting effect was assessed as inert (change from n to n), augmentative (from n to n+1) or reductive (from n to n-1).

### Thermodynamic stability

Minimum free energy (MFE) value of each sequence piece was calculated with ‘mfold’ library of Python3.7.

### Statistical analysis

Chi-square, t-test, Z-test and correlation test were performed with the ‘statsmodels’ library of Python3.7.

## Results

### Number of the common variants creating m6A motif inside G4 structure are higher than the rare variants

Since, both m6A and G4 found in pre-mRNA are known to affect splicing, localization, translational efficiency and decay of the RNAs, variants in non-spliced sequence of the selected MIM genes were evaluated in terms of their effect on m6A motif Number within putative G4 structures. While most of the variants (94,6%, 50.933 variants) were found not to relate to m6A motif (n → n; inert variants), 75% (2009 variants) of the remaining created a novel m6A motif (n → n+1; augmentative variants), and 25% (851 variants) abolished a preexisting m6A motif (n→ n-1; reductive) inside a putative G4-structure (Supplementary File 1). It was found that the Number of rare variants which create a novel m6A motif or increase the number of preexisting m6A motifs inside a G4 structure were statistically lower (p=3.02e-6) than that of the common variants (Table 1).

**Table 1:** Count of variants which decrease (n→n-1), increase (n→n+1) or does not change (n→n) the m6A motif number (n) inside or outside the G4 structure in pre-mRNA. Rare variants (MAF<0,01) causing increase of m6A motif number was significantly lower than excepted while inside the G4 structure, and higher then excepted while out of the G4 structure.

| m6A motif location | m6A motif number |  | MAF value | Variant count |  | Chi-square |
| --- | --- | --- | --- | --- | --- | --- |
|  | Reference allele | Variant allele |  | Observed | Expected |  |
| Inside G4-motif | n | n-1 | <0.01 | 509 | 529.913 | 0.82 |
|  |  |  | >=0.01 | 763 | 742.087 | 0.58 |
|  | n | n | <0.01 | 31814 | 31661.47 | 0.73 |
|  |  |  | >=0.01 | 44186 | 44338.53 | 0.52 |
|  | n | n+1 | <0.01 | 1174 | 1305.619 | 13.26 * |
|  |  |  | >=0.01 | 1960 | 1828.381 | 9.47 * |
| Outside G4-motif | n | n-1 | <0.01 | 994 | 919.87 | 5.97 |
|  |  |  | >=0.01 | 7913 | 7987.12 | 0.68 |
|  | n | n | <0.01 | 23410 | 23836.62 | 7.63 |
|  |  |  | >=0.01 | 207396 | 206969.37 | 0.87 |
|  | n | n+1 | <0.01 | 1516 | 1163.5 | 106.79 ** |
|  |  |  | >=0.01 | 9750 | 10102.49 | 12.29 ** |

\* $p=3.02e-6$ , total chi-square: 25.41, $df=2$ , \*\* $p=6.98e-30$ , total chi-square: 134.26, $df=2$

To clarify whether this situation is limited to G4-included regions, the variants which have a flanking region that is not included in any putative G4 (G4-not-included regions) were analyzed. Contrary to G4-included regions, the Number of rare augmentative variants, which creates a novel m6A motif or increases the number of preexisting m6A motifs inside G4-not-included regions, were found statistically higher (p=6.98e-30) than that of common variants (Table 1).

Since both m6A modification and G4 structures are involved in translational rate, localization and stability of mRNA, variants were evaluated in spliced RNA sequence, as well. While the difference between rare and common variants in G4-not-included regions remained statistically significant (p<0.001), there was no statistically significance in distribution of variant counts inside G4 (Table 2).

**Table 2:** Count of variants which decrease (n→n-1), increase (n→n+1) or does not change (n→n) the m6A motif number (n) inside or outside the G4 structure in mRNA. Rare variants (MAF<0,01) causing increase of m6A motif number was significantly lower than excepted while inside the G4 structure, and higher then excepted while out of the G4 structure.

| m6A motif location | m6A motif number |  | MAF value | Variant count |  | Chi-square |
| --- | --- | --- | --- | --- | --- | --- |
|  | Reference allele | Variant allele |  | Observed | Expected |  |
| Inside G4-motif | n | n-1 | <0.01 | 0 | 1.09 | 0.128 |
|  |  |  | >=0.01 | 13 | 11.9 | 1.395 |
|  | n | n | <0.01 | 71 | 68.2 | 7.995 |
|  |  |  | >=0.01 | 737 | 739.79 | 86.728 |
|  | n | n+1 | <0.01 | 1 | 2.73 | 0.320 |
|  |  |  | >=0.01 | 31 | 29.29 | 3.434 |
| Outside G4-motif | n | n-1 | <0.01 | 695 | 655.42 | 2.39 |
|  |  |  | >=0.01 | 5443 | 5482.57 | 0.28 |
|  | n | n | <0.01 | 12698 | 12857.69 | 1.98 |
|  |  |  | >=0.01 | 107714 | 107554.3 | 0.23 |
|  | n | n+1 | <0.01 | 756 | 635.88 | 22.69 * |
|  |  |  | >=0.01 | 5199 | 5319.11 | 2.71 * |

\* $p < 0.0001$ , total chi-square: 30.299, $df=2$

### Rare variants creating m6A motif inside G4 avoid being close to 3′-side and prefer to be near 5′-side of pre-mRNAs

Independently of their frequency, both inert and reductive variants displayed an equal distribution throughout pre-mRNAs. Distribution of the augmentative rare variants displayed a statistically significant shift from 3′-end to 5′-end of the pre-mRNAs (Fig 1).

**Figure 1:**
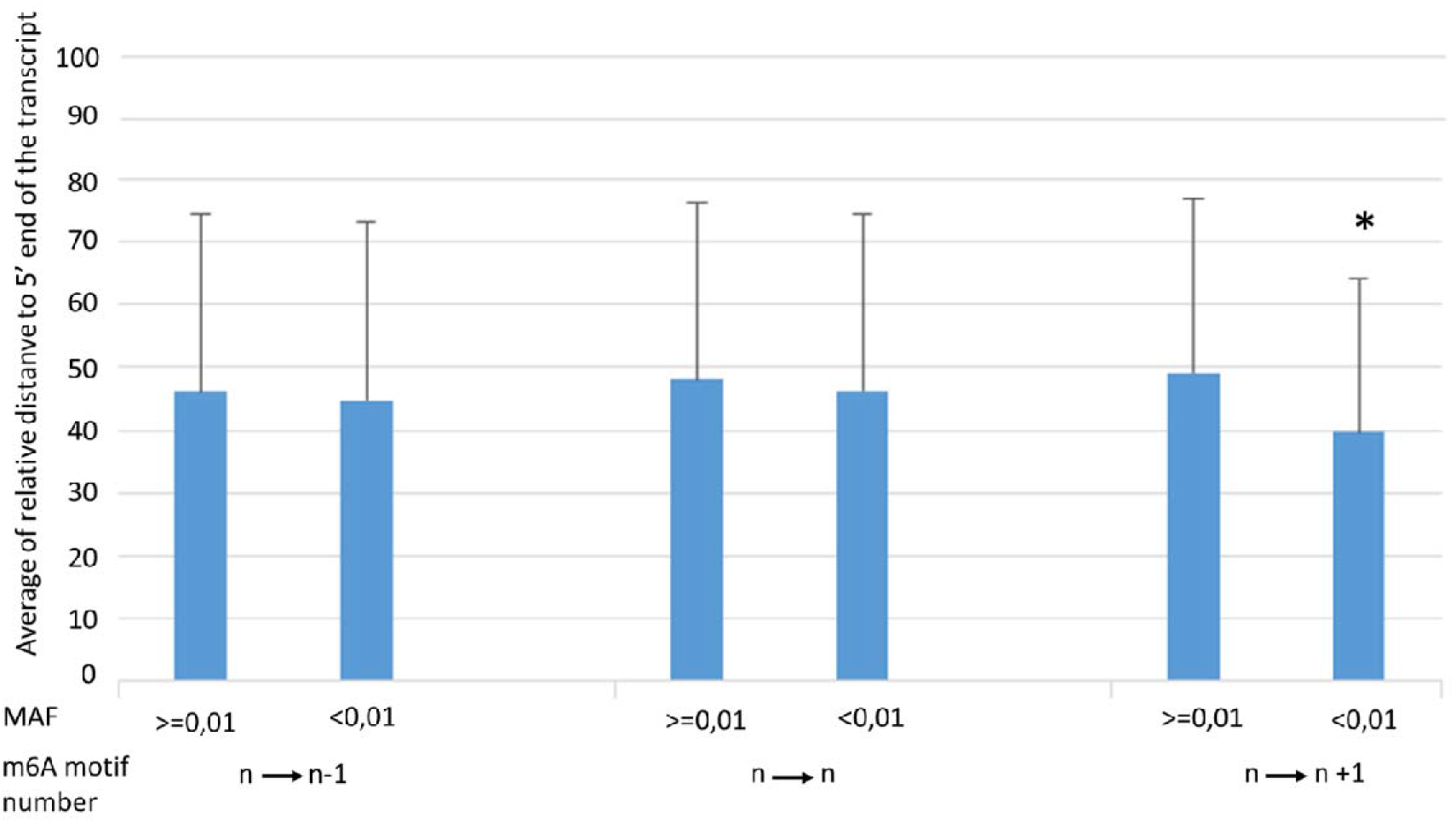
Comparison of mean distribution of common (MAF>=0.01) and rare (MAF<0.01) variants regarding their effect on m6A-motif number inside G4. Distribution of the rare variants that increase m6A number have closer location to 5′-side. (*p<0.001)

Because the total length and exon-intron contents of each transcript are different, relative position rather than absolute position of the variants is more appropriate for the comparison of their position-dependent effect in pre-mRNAs. To investigate the position-dependent effect of the variants on m6A-G4 colocalization in more detail, their relative positions were clustered as positional quartiles. Distribution of the rare variant ratio through positional quartiles revealed that ratio of inert variants displayed an equal distribution throughout pre-mRNAs. Except for the second quartile, the ratio of reductive variants also displayed an equal distribution. In the second quartile, the ratio of reductive rare variants was found significantly lower than expected. The ratio of augmentative variants displayed unequal distribution in all quartiles. Except for the last quartile, and especially in the first quartile, the ratio of augmentative rare variants was higher than expected. In the last quartile, however, it was lower than expected (Fig 2).

**Figure 2:**
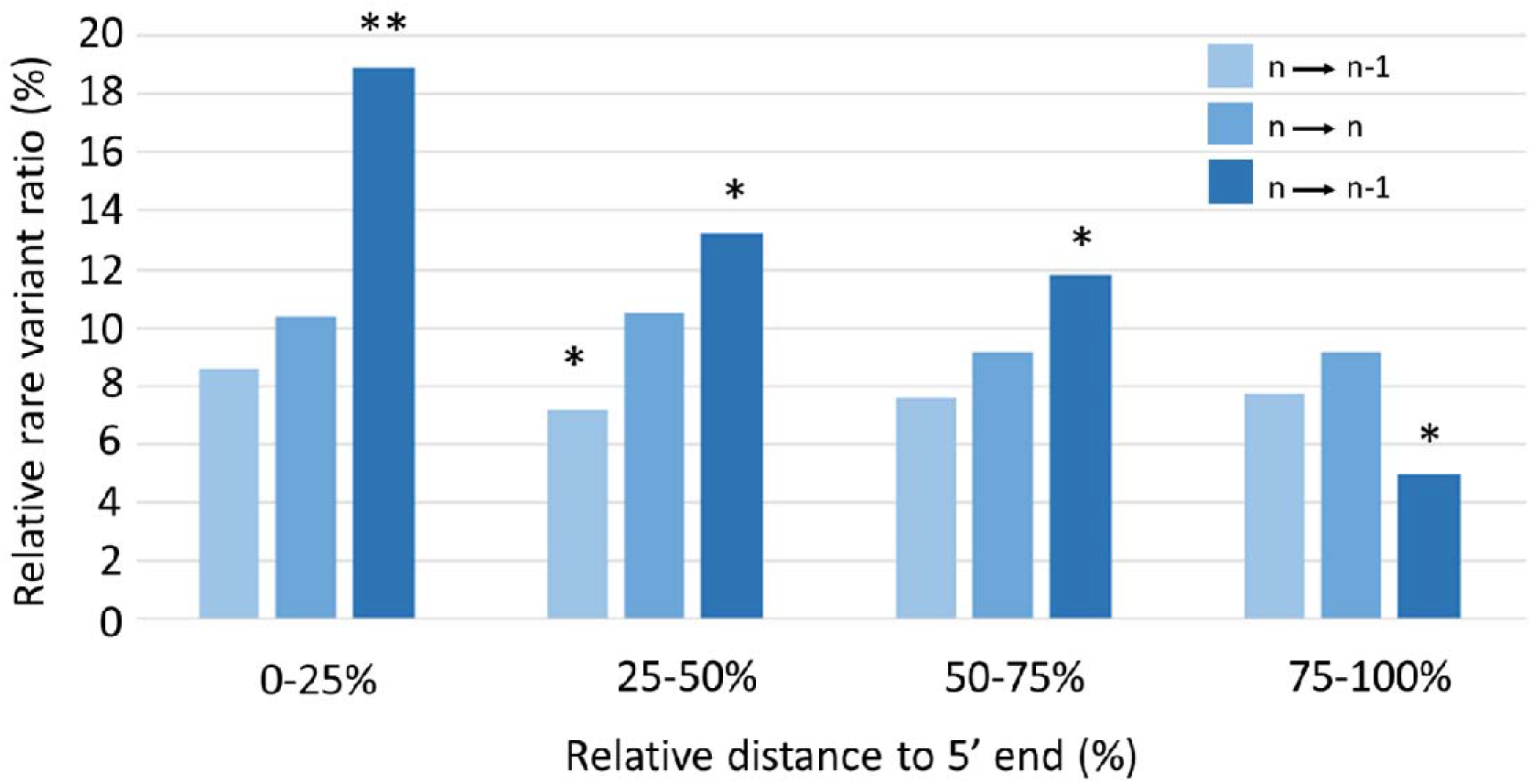
Distribution of rare variant/common variant ratio throughout relative position quartiles. The rare variants that increase m6A number display a gradually decreasing pattern from 5′- to 3′-side. (*p<0.001, **p<0.0001)

To test whether there is a difference between the distributions of variant positions throughout G4 structure, relative positions of the variants in G4 sequence were compared. There was not found any difference between over-G4 distributions of the variants (Data not shown.).

### Colocalization of m6A and G4 seems to increase the thermodynamic stability of the G4 structure

Because of the crucial involvement of thermodynamic stability in G4-structure formation, minimum free energy (MFE) values of the putative G4-structures were compared considering the variants and m6A motif numbers.

Rare variants were shown to have lower MFE value, especially in reductive variants (Fig 3a). When reference and variant bases were taken into account, it was observed that the alternative variant had lower MFE than reference allele in each group (Fig 3b). While reductive rare variants were found to have lowest MFE, augmentative common variants were found to have highest MFE value (Fig 3a). To test whether this result was due to the presence of m6A motif, MFE values of G4 sequences were compared in terms of m6A motif number. MFE values were observed to be inversely related to m6A motif count (Fig 3c). Putative G4 structures with three m6A motifs were found to have lowest MFE values (Fig 3d).

**Figure 3:**
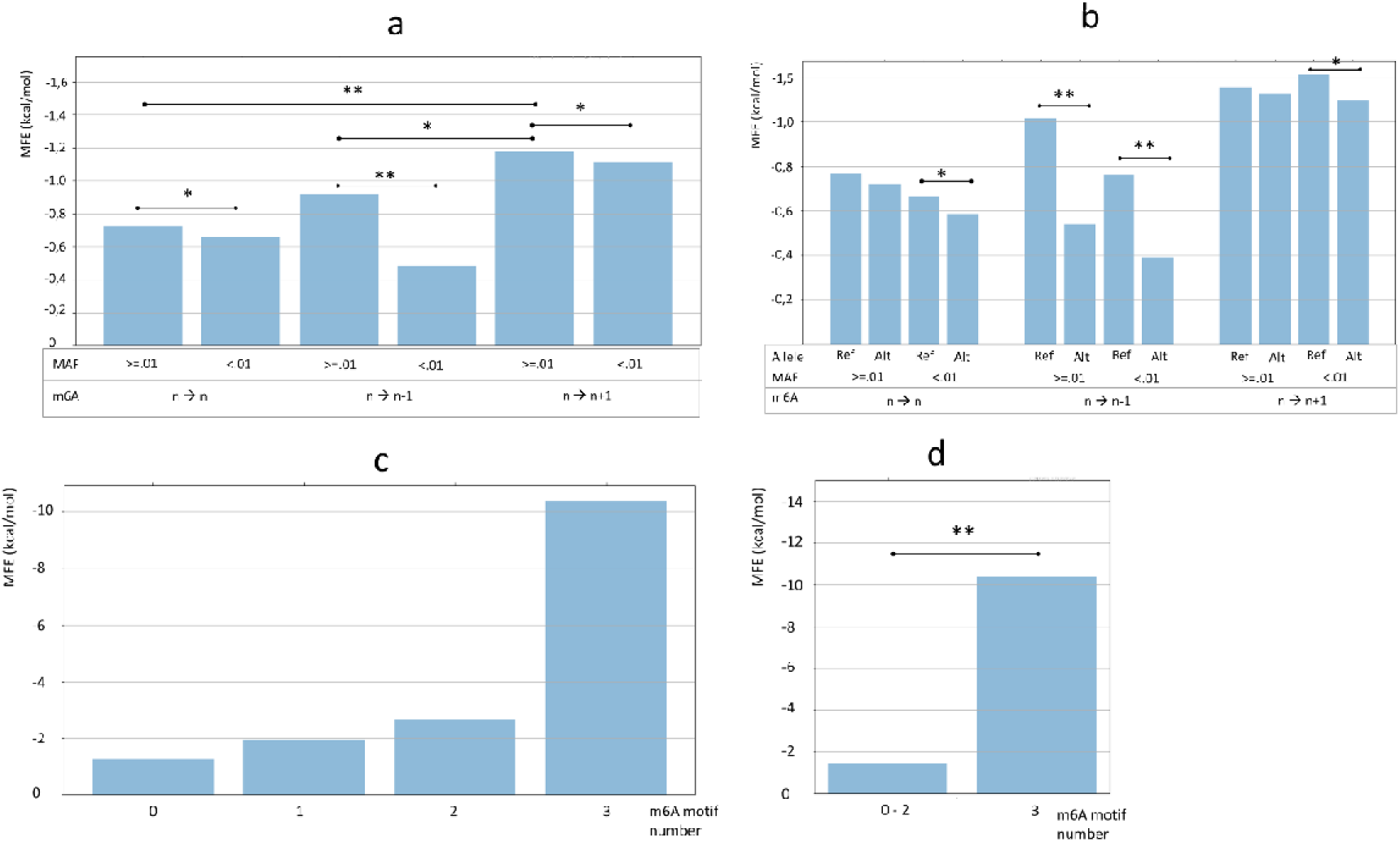
Distribution of G4-MFE values throughout relative position quartiles (a, b) and the effect of m6A number on G4-MFE value (c, d). The effect of m6A on MFE value depends on frequency (a), alleles (b) and m6A number (c), especially three m6As (d).(*p<0.001, **p<0.0001)

### Overlapping m6A-G4 may modulate G4 stability in a position-dependent manner

Due to position-dependent colocalization of m6A-G4 and m6A-dependent stability of G4, we can suppose that G4-stability may also be modulated with a position-dependent manner. Additionally, because of the relationship between overlapping m6A-G4 and allele frequency, it is possible that G4-stability may depend also on allele frequency of the variants. In order to test these possibilities MFE values of putative G4 structures were compared considering their relative position and minor allele frequency of the variants that create m6A motif.

Distribution of MFE values over positional quartiles has shown that MFE values are prone to decrease from 5′-end to 3′-end of the pre-mRNAs (Fig 4). To see whether this position-dependent decrease depends on variant alleles, MFE values were evaluated in terms of alleles. Effect of the variant alleles on MFE values was found to depend on their relative position (Fig 5a). Besides, the difference between MFE values of variant allele and reference allele also showed a dependence on relative position (Fig 5b). The MFR-difference (Variant allele MFE minus reference allele MFE) value had a decreasing pattern throughout pre-mRNA. When the distribution of MFE-difference was reevaluated considering the frequency of variant alleles, it was revealed that the decrease in MFE-difference was started closer to 5′-end with common alleles (Fig 5c), and closer to 3′-end with rare variants (Fig 5d).

**Figure 4:**
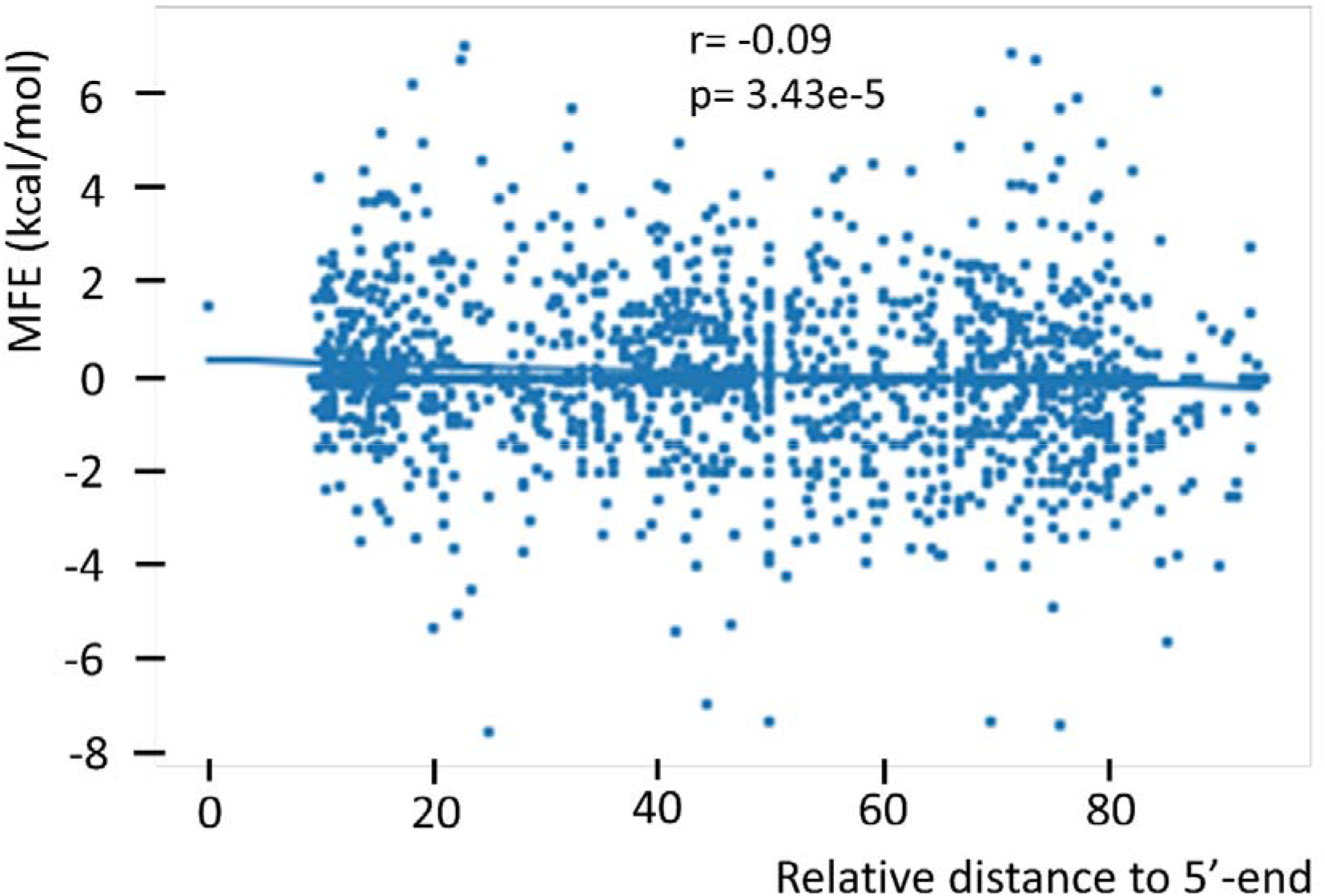
Correlation between G4-MFE values led by m6A-creating variants and their relative position. Decreasing MFE value suggests an increased stability of G4 from 5′- to 3′-end.

**Figure 5:**
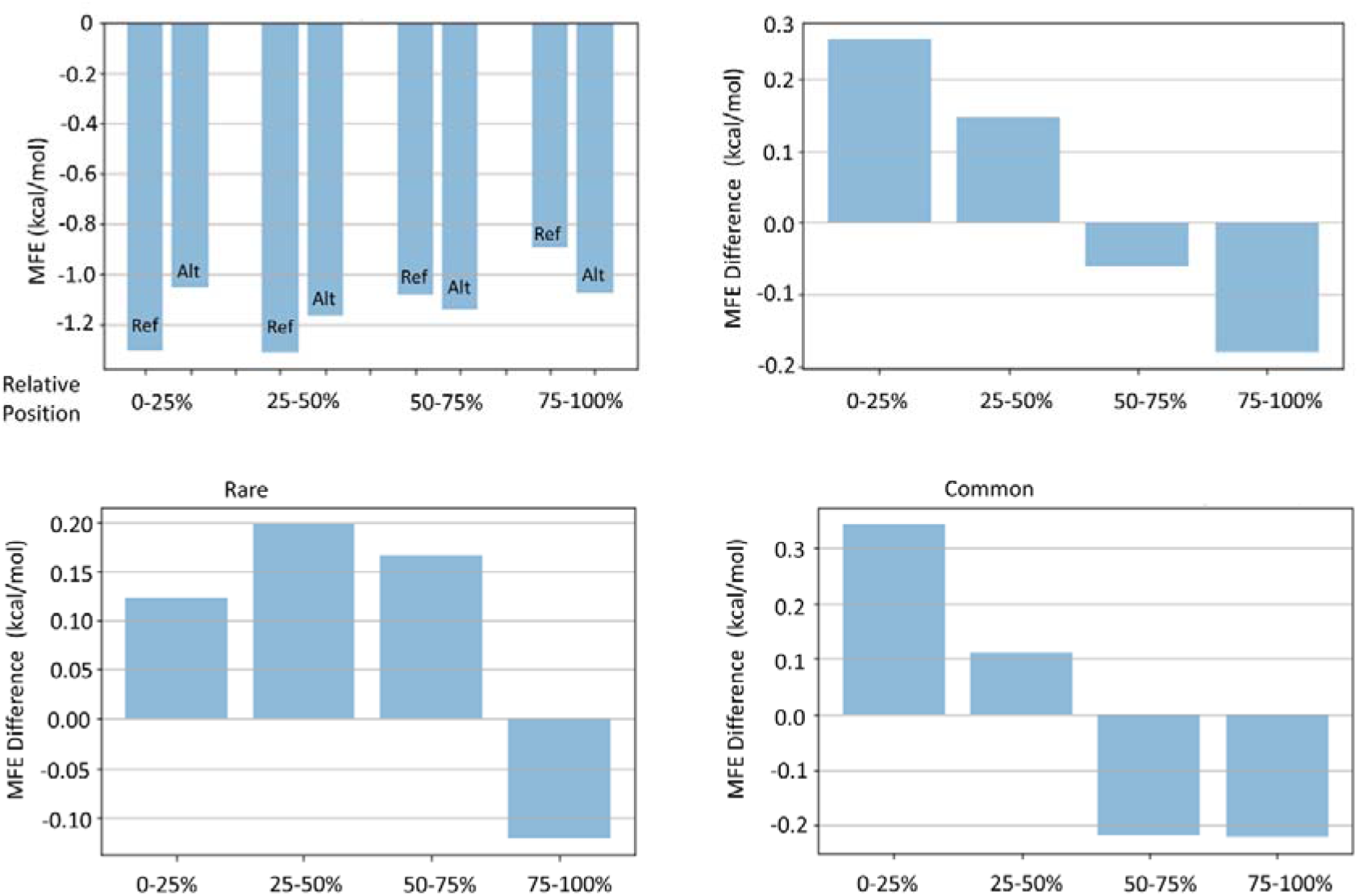
Position-dependen decrease in MFE values shown difference between reference (Ref) and alternative (Alt) allele of the variants (a). The position dependency of MFE values is more clear when MFE differences between Ref and Alt alleles are considered (b). MFE differences of common variants (c) and rare variants (d) have different distribution pattern.

## Discussion

Post-transcriptional modifications like m6A, and secondary structures like G-quadruplex, are principal actors in RNA processing. While m6A modification is controlled by specific enzymes, formation of G4 structures largely relies on physico-chemical conditions and thermodynamic rules. Nevertheless, both lead similar consequences in terms of their effects on RNA processing, but with different manners. The dependence on a consensus sequence is another common feature of m6A and G4. While m6A modification targets the adenine in the third position of DRACH ([A/G/U][A/G]AC[A/C/U]) motif [37], G4 structure is formed mostly from sequence with G_1-3_[N_1-7_G_1-3_]_3_ motif [17, 20, 21].

Besides the separate roles of m6A modification and G4 structure, their colocalization has also been shown to have functional importance. Crosstalk between m6A modification and G4 structures seems bidirectional. For instance, m6A modification was shown to affect the stability of R-loop, a G4 structure formed by DNA:RNA hybrid strands [33, 34]. Mutually, G4 structures were shown to modulate m6A modification in some viral genomes like HIV, Zika, Hepatitis B, and SV40 [36]. Supposing that a variant within G4-forming sequence may create or abolish m6A motif (DRACH), this variant can be thought to be under selective pressure depending on the functional effect of the resulting overlapping m6a-G4 status. In this respect, we can expect a correlation between frequency and colocalization ability of the variants. In this study, the supposed impact of m6A-G4 colocalization on the variant frequency was investigated. For this purpose, single nucleotide variants in selected disease-associated human genes were evaluated in terms of their allele frequency and m6A-G4 colocalization ability.

First and outstanding result of this study was that the variants creating the m6A motif inside a G4 structure are prone to have higher frequency. In contrast, such variants have lower frequency if located out of the G4 structure. Colocalization-dependent higher allele frequency of such variants suggested that overlapping m6A-G4 may have a protective role.

To understand how a variant creating m6A motif inside a G4 structure may play a protective role and gain a favorable feature, distribution of the colocalization-leading variants throughout pre-mRNAs was evaluated, since both m6A modification and G4 structure display their functional effects in a position-dependent manner [11, 38].

The results showed that the position of the colocalization-leading variants has importance for the underlying mechanism. In previous studies, it has been shown that m6A motif distribution throughout RNA molecules is not equal. Many studies presented that m6A residues were enriched in 5′ untranslated regions (UTRs), around stop codons and in 3′ UTRs adjacent to stop codons in mammalian mRNAs [5, 6, 39]. Similarly, G4 structures were also reported to be overrepresented in 5⍰- and 3⍰-UTRs [22, 40].

Findings of this study suggest that the functional consequence of the m6A-G4 colocalization may have selective pressure on the colocalization-leading variants in a position-dependent manner. While G4 structures hosting an m6A motif created by a common variant showed equal distribution throughout the pre-mRNA, G4 structures overlapping with m6A motifs created by a rare variant are prone to avoid 3′-side of the pre-mRNAs. This result might mean that m6A-G4 colocalization can be tolerated when found in the near 3′-side, but not in the central region and especially near 5′-side of the pre-mRNA. Here, we can suppose that any variant that creates a novel m6A motif inside a G4 structure may become favorable, and display an increased population frequency, if found near the 3′-side. On the other hand, the same type of variant will be unpreferable if found in the transcript body, and undesirable if found near the 5'-side of the pre-mRNA. The latter is expected to display a decreasing allele frequency. The favorability of m6A-G4 colocalization in 3′-side may be due to the supported function of the pre-existing G4 structure. For instance, as represented in Fig 6, the G4 structure supported by m6A in 5′UTR may reduce the translational efficiency of mRNA, since G4 structures in 5′UTR are known to affect cap-dependent and cap-independent translation [41]. Similarly, the G4 structure supported by overlapping m6A in 3′-side may enable the transcript to produce alternative products, through modulating alternative splicing and alternative poly-adenilation of pre-mRNA [23, 41]. It is also possible that m6A can modulate miRNA binding through stabilizing G4 structure after splicing [40].

**Figure-6:**
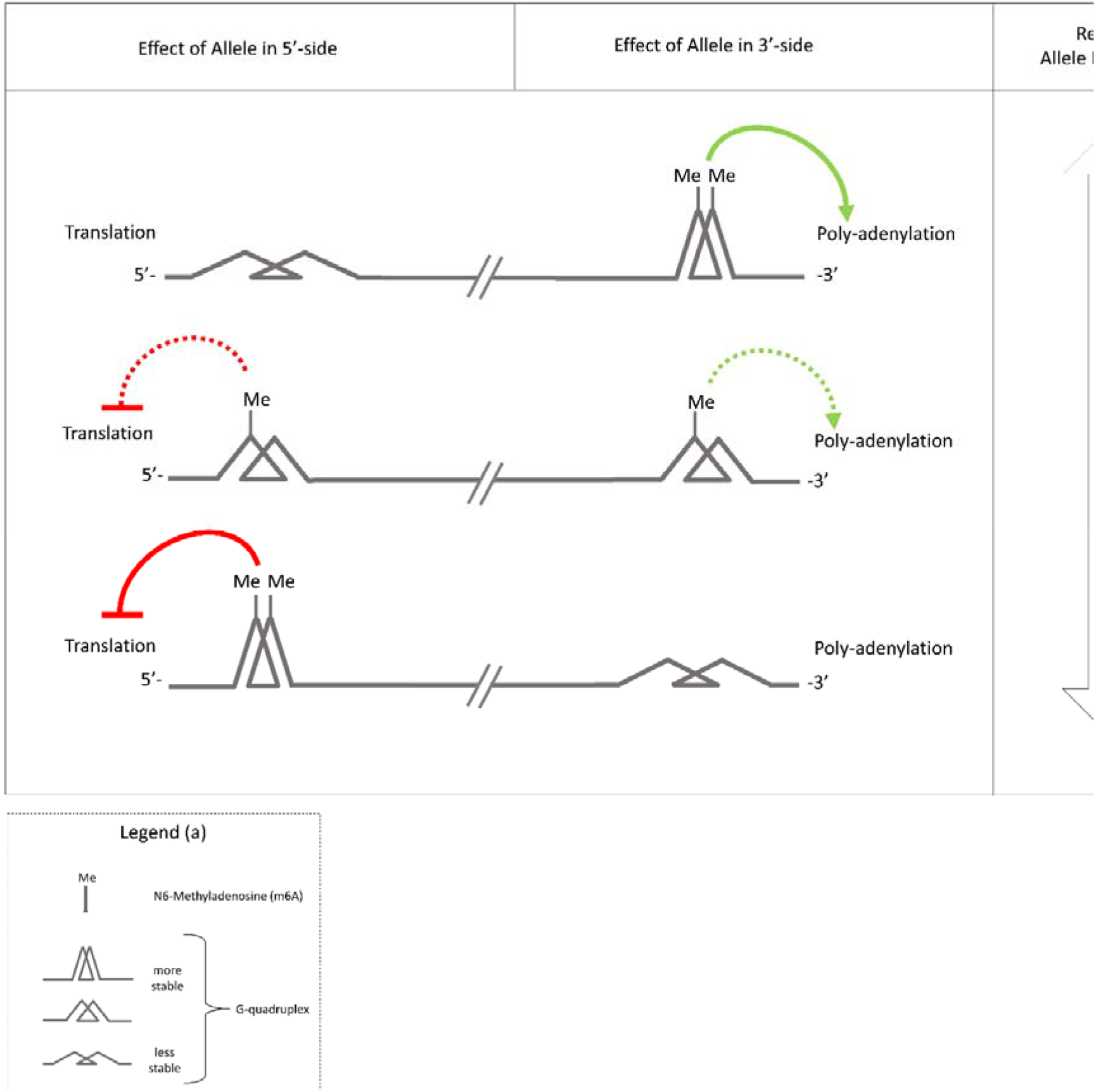
A hypothetical model to explain the position-dependent effect of m6A-G4 colocalization. A variant that creates m6A motif overlapping with G4 is prone to have higher allele freauency in 3′-side, because G4 supports poly-adenylation. In 5′-side, however, it is prone to have lower allele freauency to allow translation. Position-dependen decrease in MFE values shown difference between reference (Ref) and alternative (Alt) allele of the variants (a). The position dependency of MFE values is more clear when MFE differences between Ref and Alt alleles are considered (b). MFE differences of common variants (c) and rare variants (d) have different distribution pattern.

There may be many consequences of the m6A-G4 colocalization. Creation of novel docking sites for specific proteins can be considered in this respect. Both m6A and G4 structures are known to be recognized by specific proteins [26, 43, 44]. Therefore, colocalization of m6A modification and G4 structure may lead to competitive or cooperative interaction between these proteins. Testing these possibilities requires experimental methods dealing *in vitro* RNA-protein interactions. Another possible consequence of the m6A-G4 colocalization is the changing thermodynamic properties of the structure. Thermodynamic stability is crucial for the formation of G4 structures [45, 46]. In contrast, m6A formation is regulated by specific enzymes rather than thermodynamic status of the flanking sequence [47, 48]. However, m6A modification can affect thermodynamic stability of the RNA, as reported to marginally reduce the stability of A:U base pairing [49]. Similarly, in recent studies, colocalization of m6A and G4 was shown to alter the stability of DNA:RNA hybrid quadruplexes, known as R-loop [33, 34]. In some of these studies, m6A was reported to promote G4 folding, while in some others m6A was demonstrated to downregulate G4 formation. These contradictory findings are still discussed. Besides, G4 was also disputed to modulate m6A modification as shown in viral RNAs [34]. The overlapping m6A and G4 in 3′UTR of viral RNAs revealed that the folded G4 structures may guide the enzymatic adenine methylation in DRACH motifs [36]. Moreover, enrichment of the overlapping m6A and G4 in viral RNAs was reported to be critical for the impact of m6A on viral fitness; as shown in HIV-1 [50]. In eukaryotes, overlapping m6A and G4 was shown in 3′UTRs of mRNAs, as well [51, 52].

Based on the previous studies suggesting the synergy between m6A and G4, we can suppose that significance in the distribution of the variants leading m6A-G4 colocalization may be resulted from the changed stability of the G4 structures. To test this possibility, MFE values of G4 structures were calculated. DRACH (m6A) motifs were observed to cause a decreased MFE, which means an increased thermodynamic stability of the G4 structure. Number of m6A motifs inside the G4 structure also seems crucial for the degree of the stability.

The position-dependency of overlapping m6A-G4 may explain the mechanism responsible for difference in allele frequency among the variants leading m6A-G4 colocalization. Based on the findings of this study, we can suppose that a m6A-G4 colocalization-leading variant will prone to have higher allele frequency, if decreases G4 stability when located near 5′-side, and increases it when located near 5′-side of the pre-mRNA (Fig 7).

**Figure-7:**
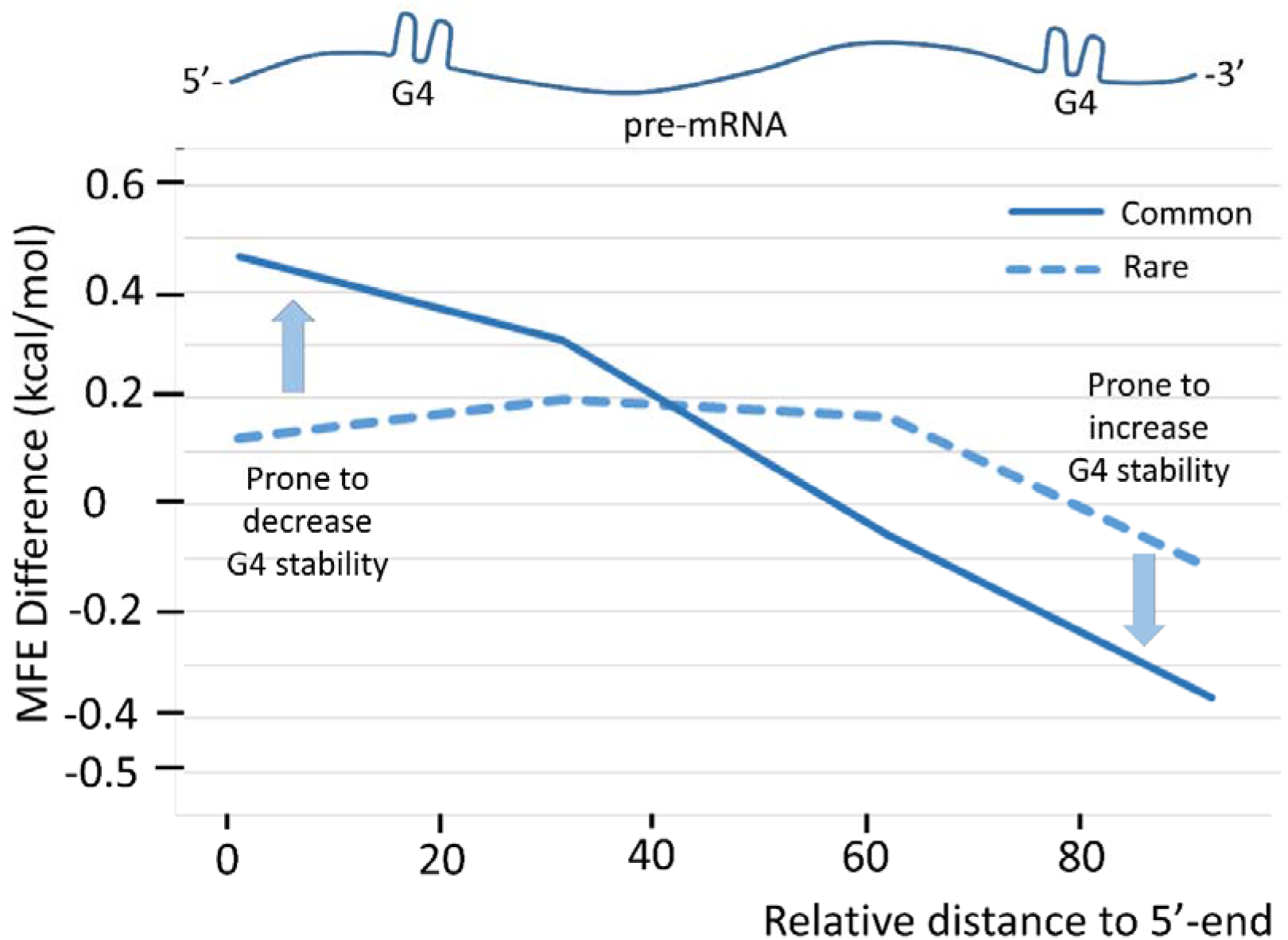
The position-dependent effect of m6A-G4 colocalization can be explained by MFE difference. The variants that create m6A motif overlapping with G4 may have better fitness due to decreased G4 stability near 3′-side and increased G4 stability near 3′-side.

The preliminary results of this study suggest that m6A overlapping with G4 structure may have functional consequences with an unknown mechanism. At least, it seems likely that this mechanism needs position-dependent stability of G4 structure.

In summary, we can conclude that the fitness, and consequently frequency of a variant creating m6A motif is prone to become higher or lower depending on whether it is located inside or outside the G4 structure, respectively. Furthermore, the frequency of these variants may depend on both their position and their effect on the thermodynamic stability of G4 structure. If located near 5′-side and destabilizes G4 or located near 3′-side and stabilizes G4, a variant creating m6A motif is prone to have higher fitness and frequency.

