## Supplemental Table for "Thermodynamic stability of G-quadruplex and overlapping m6A: Position-dependent collaborators in allele frequency and fitness?" (from another version of this preprint: version 1, 2021-04-29)

**Supplementary Table 1:** Number of rare (MAF<0.01) and common (MAF>=0.01) variants which alter m6A motif number (n) inside G4 motif in pre-mRNA

| **m6A motif number inside G4-motif** | | **MAF value** | **Variant number** | | **Chi-square** |
| --- | --- | --- | --- | --- | --- |
| **Reference allele** | **Variant allele** |  | **Observed** | **Expected** |  |
| n | n | <0.01 | 31814 | 31661.46 | 0.73 |
|  |  | >=0.01 | 44186 | 44338.53 | 0.52 |
| 0 | 1 | <0.01 | 1044 | 1138.56 | 7.85 * |
|  |  | >=0.01 | 1689 | 1594.43 | 5.6 * |
| 1 | 0 | <0.01 | 455 | 471.58 | 0.58 |
|  |  | >=0.01 | 677 | 660.41 | 0.41 |
| n | n-1 | <0.01 | 54 | 58.32 | 0.32 |
|  |  | >=0.01 | 86 | 81.67 | 0.22 |
| n | n+1 | <0.01 | 130 | 167.05 | 8.21 * |
|  |  | >=0.01 | 271 | 233.94 | 5.86 * |

* p<0.0005, total chi-square: 30.36, df=4
